# DNA Methylation in Sex Chromosomes and Genes under Selection Underpins the Emergence of Sexual Size Dimorphism

**DOI:** 10.1101/2025.04.14.648707

**Authors:** Qian Wang, Xiancai Hao, Dafni Anastasiadi, Kaiqiang Liu, Shuo Li, Bo Feng, Rui Wang, Michael G. Ritchie, Yuyan Liu, Hong-Yan Wang, Lili Tang, Francesc Piferrer, Changwei Shao

## Abstract

Sexual size dimorphism (SSD) refers to the differences in body size between males and females. Epigenetics such as methylation contribute to shaping phenotypes, nevertheless, their role in SSD is still unclear. Here, we used *Cynoglossus semilaevis*, a species with larger females known as female-biased SSD, as a model to investigate the role of methylation in the emergence and regulation of SSD. Methylomes and transcriptomes were constructed in four tissues (brain, liver, muscle, and gonad) from three key life stages: juveniles (3-months post fertilization), male mature stage (1-year post fertilization; ypf), and female mature stage (3 ypf). In 1 ypf and 3 ypf females showed lower methylation, while the number of differentially methylated regions between sexes increased during the lifetime. Genes at the top of the growth hormone/insulin-like growth factor (GH/IGF) axis and hypothalamus-pituitary-gonadal axis (*gh1*, *igf1*, *gnrhr2r-like*, *err*γ, *etc*.) showed differential methylation between sexes, consistent with sex-related differences in growth. The female-specific W chromosome showed higher methylation than the autosomes or the Z chromosome. Genes of the IGF signaling network that negatively regulate growth and located on W chromosome were hyper-methylated than their Z homologs. Furthermore, genes with a faster evolutionary rate in *C. semilaevis* exhibited lower methylation levels than the background genes, suggesting an important evolutionary role for DNA methylation in shaping SSD. Our results provide a comprehensive depiction of methylation regulation of SSD in vertebrates, and improve our understanding of how methylation regulation can help organisms to respond to natural selection, and allocate resources between the sexes.

## Introduction

Sexual dimorphism is the divergence of traits exhibited between females and males in the same species, including secondary sex characteristics, growth, behavior, immune function, aging, *etc.* [1,2]. This phenomenon occurs widely in the animal kingdom and has captivated the attention of many biologists. Sexual size dimorphism (SSD), which refers to differences in body size between adult males and females, is one of the most obvious traits of sexual dimorphism.

As illustrated by life history theory, the general features of individual such as growth, reproduction, and lifespan are constrained by trade-offs. Thus, they are limited by the finite resources and must compete in a way that maximizes fitness [3,4]. Therefore, when looking into SSD, aspects related to reproduction and lifetime should be considered as well as the growth trait (body size).

Hormones play an important role in regulating a vast array of physiological functions including growth and reproduction in living organisms [5−7]. Specifically, the brain-pituitary-liver axis is essential in regulating somatic growth and development. It consists of somatostatin and growth hormone (GH) releasing hormone secreted by hypothalamus, GH secreted by pituitary, GH receptor and insulin-like growth factor (IGF) secreted by liver [8]. Disturbances in the GH-IGF network may cause failure to show sufficient catch-up growth in children with small for gestational age (SGA) syndrome [9]. On the other hand, the hypothalamus-pituitary-gonadal (HPG) axis plays an important role in regulating reproduction processes. Abnormal state of hormones, receptors, and enzymes in this axis may affect fertility or gonadal development in humans [10,11]. The molecular mechanisms underlying sexual dimorphism have been elucidated from gene expression, genetic component, and methylation regulation aspects, mainly regarding the traits of morphology, aging, and immune system between sexes [12−14]. However, such studies in relation to SSD are still limited.

The phenomenon of SSD is widespread in teleosts, characterized by their extraordinary diversity [15], which may reflect the divergent selective pressures on different sexes across taxa, such as sexual selection acting on males or fecundity selection on females [16−19]. Sexual selection favors male-biased SSD and is considered to be driven by competition over mates for reproductive success [20,21]. Fecundity selection favors female-biased SSD, since larger abdominal volume could increase fecundity [22]. Thus, natural selection has showed how environment shapes life history and phenotypic adaptations at individual level. In teleost, at least 118 fish species with SSD in the adult stage have been reported [16], a number that is likely to increase as more species are examined. Based on these 118 species, it has been found that, on average, males are somewhat smaller than females, although many exceptions do exist [23]. Thus, males are usually larger than females where male competition is present, including tilapia (*Oreochromis* spp.) and channel catfish (*Ictalurus punctatus*). Conversely, females are larger than males in species where there is increasing fecundity selection, such as sea bass (*Dicentrarchus labrax*), turbot (*Scophthalmus maximus*), and Chinese tongue sole (*Cynoglossus semilaevis*) [23,24]. The mode of reproduction and the genomic architecture of fish species make them ideal models to study the regulatory mechanism of SSD. Many studies have illustrated that sex-biased gene expression is related to SSD traits [25,26]. Yet the mechanisms regulating sex-biased gene expression are worth further exploration.

Recent research has revealed that genes shared between sexes can be differentially regulated by methylation modifiers thus create a sex-specific phenotype. For example, differential cytosine methylation in muscle may contribute to sex-specific growth phenotypes in tilapia (*O. niloticus*) [27,28]. Differences in cytosine methylation of *myogenin* have been linked to thermal plasticity of growth in Atlantic salmon (*Salmo salar*) and Senegalese sole (*Solea senegalensis*) [29,30]. Furthermore, in *C. semilaevis*, where females achieve final body sizes two to four times larger than males, approximately 14% of genetic females can sex-reverse into phenotypic males, called pseudomales, which end up in a body size similar to that of regular genetic males [31−33]. Previous studies showed that methylation modifications in high-temperature induced pseudomale *C. semilaevis* was globally inherited by their ZW offspring, which can naturally develop into pseudomales without temperature incubation [34]. These results suggested that DNA methylation could be directly involved in the phenomenon of SSD.

In the present study, we used *C. semilaevis* as a model to investigate the methylation mechanisms underlying SSD. Four important tissues (whole brain, liver, muscle, and gonad) related to growth and reproduction in three different life stages: 3-months post fertilization (mpf) (juvenile stage that sex-determination program is accomplished), 1-year post fertilization (ypf) (male sexually mature stage), and 3 ypf (female sexually mature stage) were selected from females and males. Fish were routinely reared in filtered seawater at 22°C to avoid the effect of the temperature on sex determination. High throughput sequencing of the methylome at single-base resolution and the transcriptome was performed. Our results provide a detailed molecular picture of methyl-regulated functions in SSD in vertebrates, which is helpful to understand how methylation regulation can help organisms to allocate resources between different sexes to maximize individual fitness, and to cope with the pressure of natural selection.

## Results

### Sex-dimorphic growth in *C. semilaevis*

To quantitatively characterize SSD in *C. semilaevis*, we measured growth parameters of males and females under the routine 22°C rearing condition at three developmental stages. At 3 mpf, no significant differences were observed between sexes in body length or weight. At 1 ypf, the females experienced rapid growth and became significantly larger than males (*P* < 0.001). At 3 ypf, sex-related differences were even greater (*P* < 0.001) (**Figure 1A**, Figure S1A).

**Figure 1.**
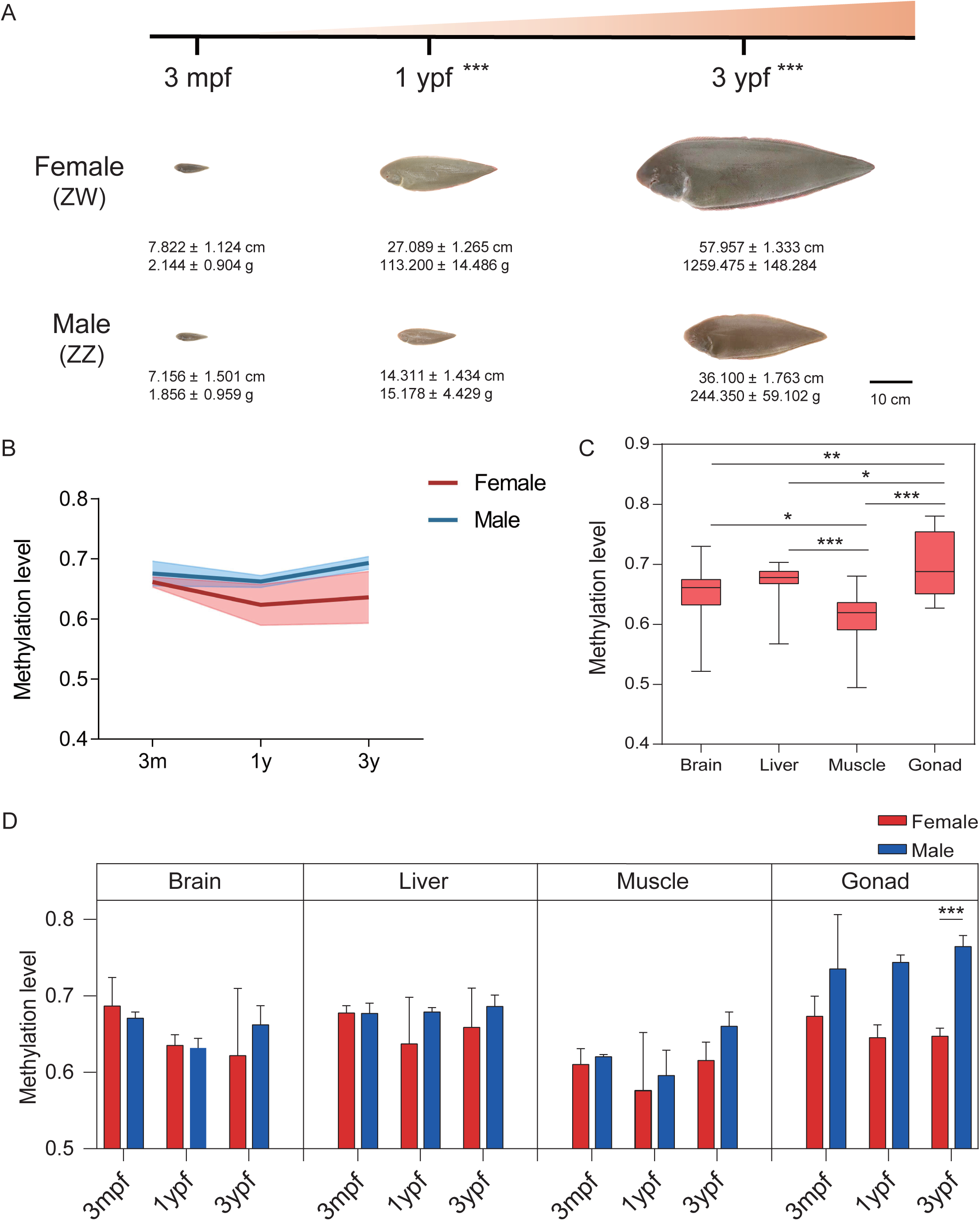
SSD in *C. semilaevis* and integrated analysis of the genome-wide DNA methylation profiles **A**. SSD in *C. semilaevis*. At 3 mpf, the average body length and body weight of *C. semilaevis* showed no statistically significant difference between females and males. At 1 ypf, the growth characteristics in females became significantly larger than males. At 3 ypf, sex-related differences were more obvious (*n* = 9). **B**. Methylation levels of mCpGs in females (red) and males (blue) from 3 mpf to 3 ypf. **C**. Overall methylation levels of the four different tissues shown by boxplots. **D**. Methylation level of gene body with promoter region in each sample shown by boxplots. Black line indicates the median, lower and upper edges stand for the 25th and 75th percentiles. Red represents female and blue represents male. SSD, sexual size dimorphism; mpf, months post fertilization; ypf, years post fertilization. *, *P* < 0.05, **, *P* < 0.01, ***, *P* < 0.001 in each comparison, *t*-test.

### DNA methylation profile in *C. semilaevis*

To explore the possible methylation mechanism underlying SSD, we generated whole genome bisulfite sequencing (WGBS) data from 72 samples of female and male *C. semilaevis* brain, liver, muscle, and gonad in 3 mpf, 1 ypf and 3 ypf, respectively. A total of 1.42 Tb methylome raw data was obtained. After data filtering, an average of 17.60 Gb clean bases per library was retained (Table S1). An average of 64.02% non-duplicated alignments were obtained, which produced an average depth of 29.24 × per strand and an average 86.25% C sites coverage for each group (Table S2).

We identified an average of 10.10 million methylated cytosines (mCs) per library, representing 13.87% of cytosines in the reference genome. Cytosines can be classified into three sequence contexts (CG, CHG and CHH where H is a nucleotide other than G). As in other vertebrates, DNA methylation mainly occurred in the CG context (90.69% of mCs), with substantially lower levels in CHG (1.99%) and CHH (7.31%) contexts (Figure S1B, Table S3). Thus, we focused on the CpG sites for subsequent analyses. Across various genomic elements, the promoter region was relatively hypo-methylated with an average methylation level of 35%, while exons and introns were hyper-methylated with average methylation level of 74% and 70%, respectively (Figure S1C, Table S4). Also, we observed a dramatic drop in methylation level around the transcription start site (TSS) (Figure S1D).

### Sexually dimorphic DNA methylation varies during development

To identify major sources of variation in WGBS data, principal component analysis (PCA) was performed. Tissue-specific difference in methylation levels dominated over differences across developmental stages or sexes (Figure S2A). We next detected overall changes of CpG methylation level (taking all tissues together) along with development stages. At 3 mpf, methylation levels were similar between sexes (females: 0.6617, males: 0.6756). At 1 ypf, methylation levels decreased in females but maintained in males (females: 0.6235, males: 0.6624). At 3 ypf, methylation divergence was accelerated (females: 0.6358, males: 0.6931) (Figure 1B, Table S5). This progressive methylation sexual dimorphism mirrored phenotypic growth differences (body weight and body length) and extended to repetitive elements (SINEs, LINEs) and regulatory regions (CpG islands, promoters) (Figure S2B).

We then quantified tissue-specific methylation levels across brain, liver, muscle, and gonads. These tissues exhibited different methylation levels, with the highest in 3 ypf male gonad (0.7643) and lowest in 3 ypf female muscle (0.5763) (Figure 1C, Table S5). Changes of methylation patterns in tissues were different between sexes and developmental stages. In general, the sex difference was not significant in all but the gonad tissues. In brain, methylation levels gradually decreased in females, while they decreased at 1 ypf and then increased at 3 ypf in males. In liver, methylation levels decreased at 1 ypf and then increased at 3 ypf in females, while showed only subtle changes in males. In muscle, methylation levels decreased at 1 ypf, then increased at 3 ypf in both sexes, with males showing higher methylation levels than females. In gonad, the methylation levels gradually decreased from 0.6730 to 0.6472 in females, while they gradually increased from 0.7350 to 0.7643 in males (Figure 1D, Table S5).

### Female and male methylome signatures during development

We then investigated the regulatory role of DNA methylation dynamics in SSD across developmental stages. While the number of differentially methylated regions (DMRs) between females and males increased during development in brain, muscle, and gonad, DMR number of liver was peaking at 1 ypf (15,440) compared to 3 mpf (11,364) and 3 ypf (14,513) (**Figure 2A**). At 3 mpf, the number of female hyper- and hypo-methylated DMRs was similar in liver (5707 vs 5657) and muscle (7800 vs 9658), whereas brain (8563 vs 4859) and gonad (4558 vs 41,431) exhibited marked imbalances. Notably, from 1 ypf on, there was an obvious increasing trend in the number of female hypo-methylated DMRs in all tissues, with liver uniquely showing the highest number of DMRs at 1 ypf (12,846) (Figure S2C).

**Figure 2.**
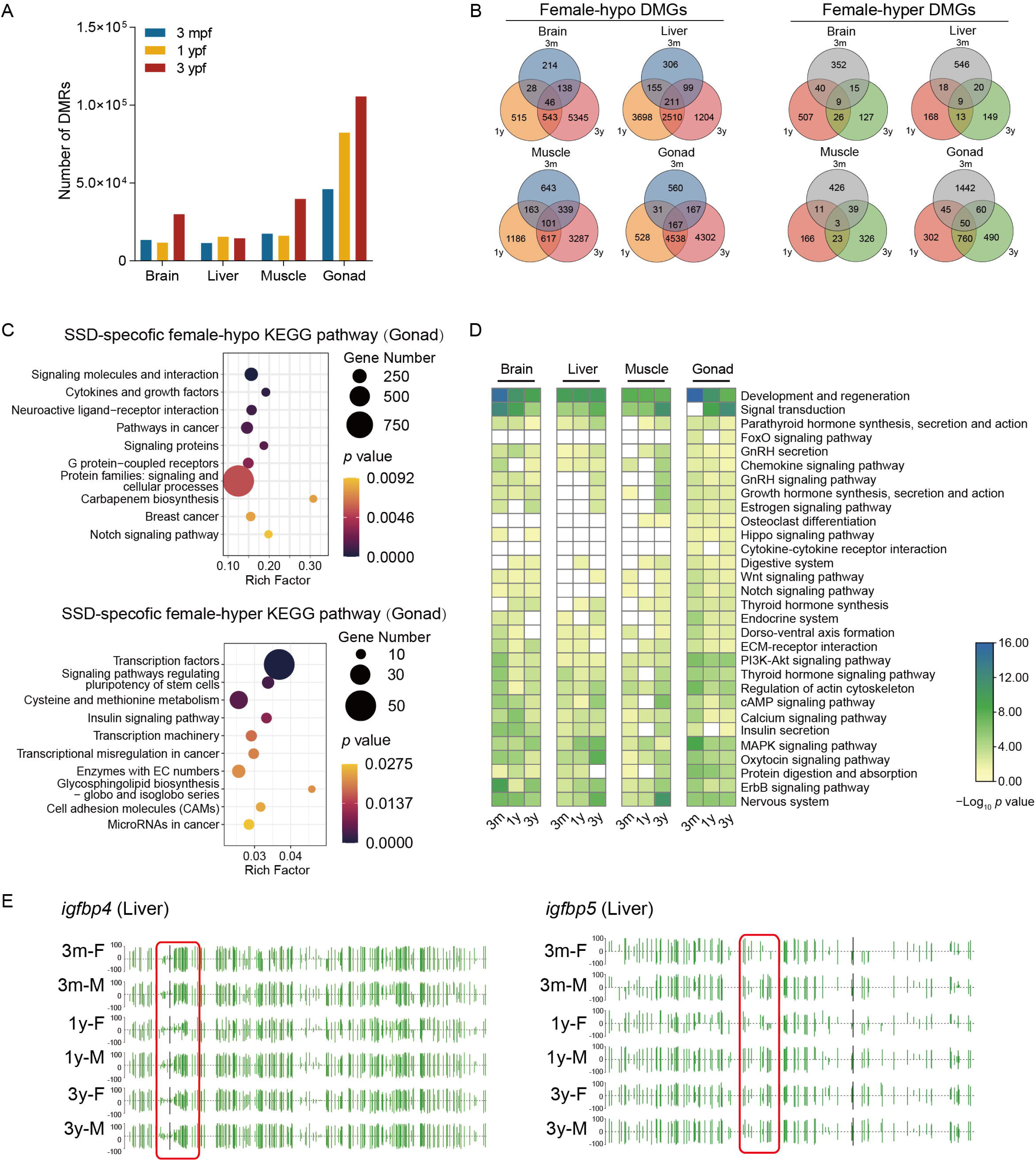
Analysis of the DMRs between female and male *C. semilaevis* **A.** Number of DMRs between sexes in different developmental stages in four tissues. **B.** Venn diagrams of SSD-specific female hypo-methylated (left) and hyper-methylated (right) DMGs in four tissues. **C**. KEGG enrichment analysis of SSD-specific female hypo-methylated (up) and hyper-methylated (down) DMGs in gonad. **D**. KEGG enrichment analysis of DMGs between sexes in four tissues. **E**. The DNA methylation level of *igfbp4* and *igfbp5* promoters shown by genome browser tracks. Green vertical lines indicate the methylation level of cytosines. The number “100” and “-100” represented the methylation level in sense and antisense strand, respectively. DMR, differentially methylated region; DMG, differentially methylated gene.

To identify the corresponding differentially methylated genes (DMGs), we screened the DMRs overlapped with gene promoter regions. Female hypo-methylated DMGs were significantly more than hyper-methylated DMGs in all tissues. There were 6829 hypo- and 1076 hyper-methylated DMGs in brain, 8183 hypo- and 923 hyper-methylated DMGs in liver, 6336 hypo- and 994 hyper-methylated DMGs in muscle, 10,293 hypo- and 3149 hyper-methylated DMGs in gonad (Figure 2B). According to the phenotypic progression of SSD in *C. semilaevis*, the SSD-specific DMGs were defined as (1) no significant differences at 3 mpf, (2) emergence of significant differences at 1 ypf, and (3) amplified differences at 3 ypf. This definition resulted 335, 401, 224, and 2494 SSD-specific female hypo-methylated DMGs, as well as 5, 6, 14, and 303 SSD-specific female hyper-methylated DMGs in the brain, liver, muscle and gonad, respectively (Figure S3,). Gonad exhibited the highest numbers of SSD-specific DMGs, suggesting its central role in SSD regulation. Kyoto Encyclopedia of Genes and Genome (KEGG) pathways enrichment analysis found female hypo-methylated DMGs significantly enriched in pathways including “signaling molecules and interaction”, “cytokines and growth factors”, and “neuroactive ligand-receptor interaction”, and female hyper-methylated DMGs significantly enriched in pathways including “transcription factors”, “signaling pathways regulating pluripotency of stem cells”, and “cysteine and methionine metabolism” (Figure 2C). Pathways of other tissues showed a tissue-specific pattern, although “protein families: signaling and cellular processes” and “G protein-coupled receptors” were found common to brain, muscle, and gonad in female hypo-methylated DMGs, while only “insulin signaling pathway” was common to liver and gonad in female hyper-methylated DMGs (Figure S4).

When taking all the DMGs between sexes across all different stages and tissues into consideration, KEGG pathway analysis showed specificity among stages and tissues. “development and regeneration” pathway was significantly enriched with time in all tissues, especially was most enriched at 3 mpf in brain and gonad. “signal transduction” pathway was gradually significantly enriched with time in all tissues, except in brain that was most enriched at 3 mpf (Figure 2D). Notably, pathways of the “GnRH secretion”, “GnRH signaling pathway”, and “growth hormone synthesis, secretion and action” were significantly enriched from 3 mpf on in the brain and gonad, which may reflect the beginning of regulation via DNA methylation prior to the appearance of phenotypic differences. Accordingly, genes encoding insulin-like growth factor-binding proteins, *i.e.*, *igfbp4* and *igfbp5*, which regulate growth via binding to IGF 1, exhibited similar methylation level between 3mpf (*igfbp4*: 0.5058, *igfbp5*: 0.3846) and 1ypf (*igfbp4*: 0.5346, 0.4279) in females, while they were hyper-methylated at 3mpf (*igfbp4*: 0.4975, *igfbp5*: 0.3645) than 1ypf (*igfbp4*: 0.5973, *igfbp5*: 0.5092) in males (Figure 2E).

### Gene expression changes across the lifetime

To investigate genes that differentially expressed between sexes and their potential regulatory association with DNA methylation dynamics, we employed RNA-seq to profile female and male transcriptomes from samples at 3 mpf, 1 ypf, and 3 ypf in brain, liver, muscle, and gonad. Genome-wide correlation analyses between methylation levels and transcriptional activity revealed a negative correlation between promoter methylation and transcription across all tissues and developmental stages (Figure S5). There was only a small number of differentially expressed genes (DEGs) between females and males at 3 mpf. Numbers of DEGs increased at 1 ypf and then slightly reduced at 3 ypf in brain, liver, and muscle. In the gonads, the number of DEGs gradually increased from 3 mpf to 3 ypf, with this increase being greater than in the other three tissues (**Figure 3A**). This was consistent with the pattern of methylation data, and resulted in the gonad being the most morphologically heterogeneous tissue in adult *C. semilaevis*. Similarly, SSD-specific DEGs were defined according to DMGs with their fold change differences. This resulted 49, 32, 34, and 2094 SSD-specific female up-regulated DEGs, as well as 12, 13, 28, and 3922 female down-regulated DEGs in the brain, liver, muscle and gonad, respectively (Figure S6). Overlapping female hypo-methylated SSD-specific DMGs vs. up-regulated DEGs and female hyper-methylated SSD-specific DMGs vs. down-regulated DEGs identified 225 and 71 genes in gonad (Figure 3B). No overlaps were detected in other tissues, with only two functionally irrelevant overlaps identified in brain and liver, respectively. KEGG analysis revealed the “ribosome biogenesis” and “DNA replication” pathways were enriched in the gonadal SSD-specific female hypo-methylated/up-regulated genes, while “alpha-linolenic acid metabolism” and “cell adhesion molecules (CAMs)” pathways were enriched in the gonadal female hyper-methylated/down-regulated genes, which may reflect the enhanced protein synthesis and cell cycle progression capacity during female rapid growth (Figure 3C).

**Figure 3.**
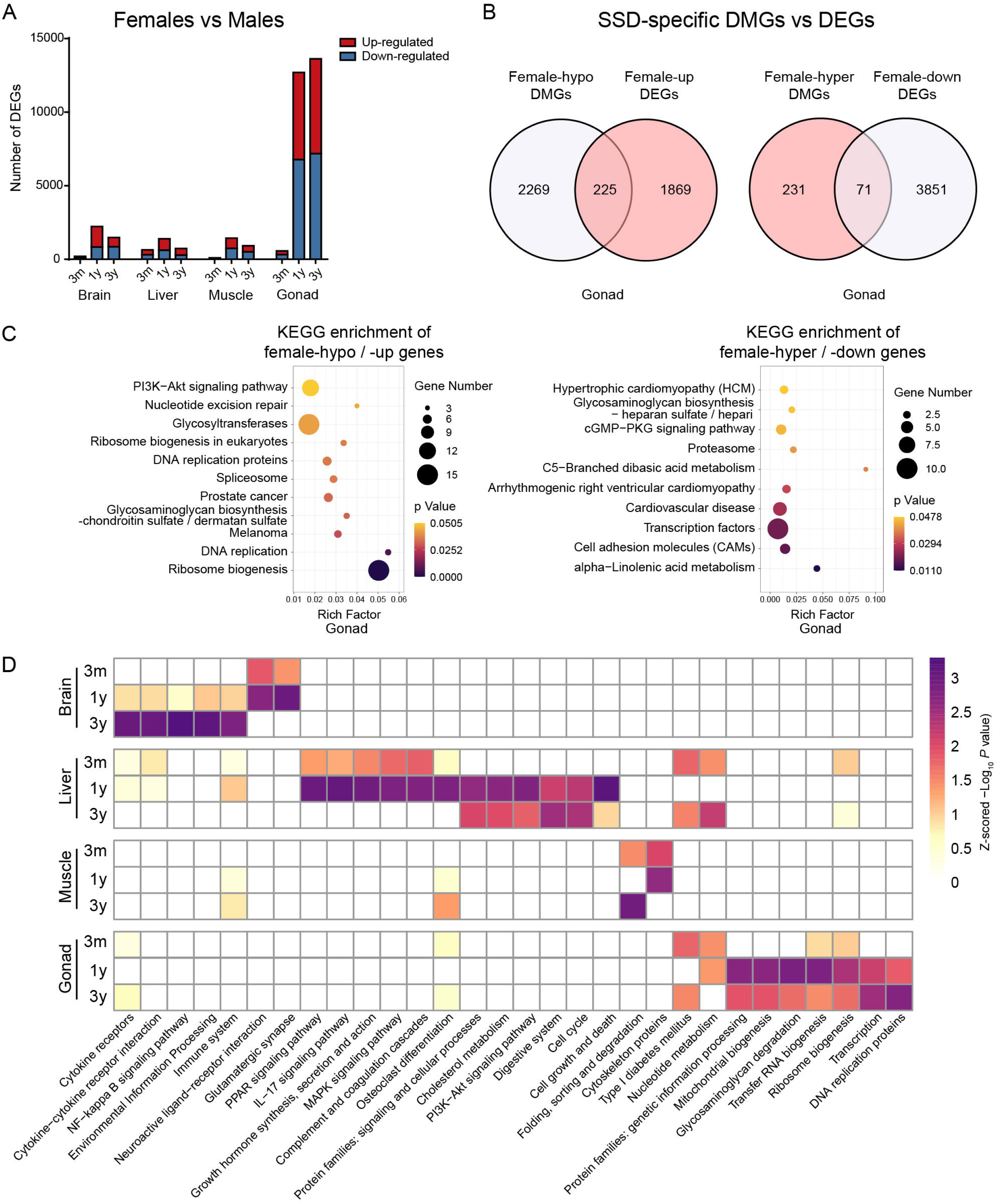
Transcriptome analysis of DEGs between female and male *C. semilaevis* **A**. Number of female up-regulated (red) and down-regulated (blue) DEGs between sexes in different stages in four tissues. **B**. Venn diagram of female hypo-methylated SSD-specific DMGs vs. up-regulated DEGs (left), and female hyper-methylated SSD-specific DMGs vs. down-regulated DEGs (right) in gonad. **C**. KEGG enrichment of SSD-specific female hypo-methylated/up-regulated (left) and hyper-methylated/down-regulated genes (right) in gonad. **D**. KEGG enrichment analysis of DMGs between sexes in four tissues. The −Log_10_ *P* value was treated by Z-score normalization. DEG, differentially expressed genes.

Taking the DMGs between sexes across all different stages and tissues into consideration, KEGG analysis revealed “cell cycle” and “cell growth and death” pathways were enriched in liver, “osteoclast differentiation” pathway was enriched in liver and muscle, “mitochondrial biogenesis” and “ribosome biogenesis in eukaryotes” were enriched in gonad, both at 1 ypf and 3 ypf, which may reflect the energy and material needed during the rapid growth. Besides, the “cytokines and growth factors” pathway was specifically enriched at 3 mpf in liver and gonad, which may reflect preparation for subsequent rapid growth of the organism. The “cytokine-cytokine receptor interaction” and “environmental information processing” pathways were increasingly enriched from 1 ypf to 3 ypf, which may be consistent with the physiological stage in which growth tends to be stable and reproduction becomes the main task (Figure 3D).

### Genes participating in growth- and reproduction-associated networks were regulated by methylation during lifetime

To evaluate the effect of methylation on growth-/reproduction-related genes, we constructed a list of 205 genes mainly located on the growth-related GH/IGF network and reproduction-related HPG axis based on the literature, and investigated their promoter methylation status in the corresponding distributed tissues (Figure S7, Table S6). These genes showed a negative correlation between their methylation and gene expression levels (*R* = −0.30, *p* < 0.001, Spearman) (Figure S8), and many of them exhibited methylation differences between females and males (**Figure 4A**, **B**). At 3 mpf, IGF receptor gene *igf1r1* (XM_008323538.3), IGF binding protein gene *igfbp2* (XM_008325023.3), myostatin inhibiting gene *fst* (XM_008336107.3) were hypo-methylated in females, indicating these growth-promoting genes might be critical upstream regulators before the females and males showed body size difference. At 1 ypf, key growth-promoting genes including growth hormone gene *gh1* (NM_001294211.1) and insulin-like growth factor gene *igf1* (NM_001294198.1) were significantly decreased in females [*gh1*: female = 0.86 (3mpf), 0.62 (1ypf), 0.57 (3ypf); male = 0.80 (3mpf), 0.71 (1ypf), 0.79 (3ypf). *igf1*: female = 0.40 (3mpf), 0.32 (1ypf), 0.42 (3ypf); male = 0.44 (3mpf), 0.41 (1ypf), 0.41 (3ypf)], with pronounced increased mRNA expression at the same stages (*P* < 0.01) (Figure 4C). The IGF binding protein gene *igfbp5* (XM_008329025.3) were also hypo-methylated in females at 1 ypf (female = 0.43, male = 0.51) and 3 ypf (female = 0.35, male = 0.51). On the other hand, growth-inhibiting somatostatin receptor gene *sstr5* (XM_008329192.3) showed a significantly increased methylation level in females [female = 0.58 (3mpf), 0.70 (1ypf), 0.60 (3ypf); male = 0.51 (3mpf), 0.49 (1ypf), 0.62 (3ypf)], with its mRNA expression level lower in females than in males (Figure 4C). Many genes involved in the IGF-Akt/PKB pathway regulating muscle growth such as *foxo3-like* (XM_008310881.3) and *mstn* (XM_008328997.3) did not show significant methylation differences between females and males. Nevertheless, their mRNA expression levels were up-regulated in females than in males (*P* < 0.05). The muscle growth promoting gene *eif2b3* (XM_008333984.2) was hypo-methylated in females at 1 ypf (female = 0.20, male = 0.27) and 3 ypf (female = 0.25, male = 0.34), with higher mRNA expression level (*P* < 0.01) (Figure 4B, C).

**Figure 4.**
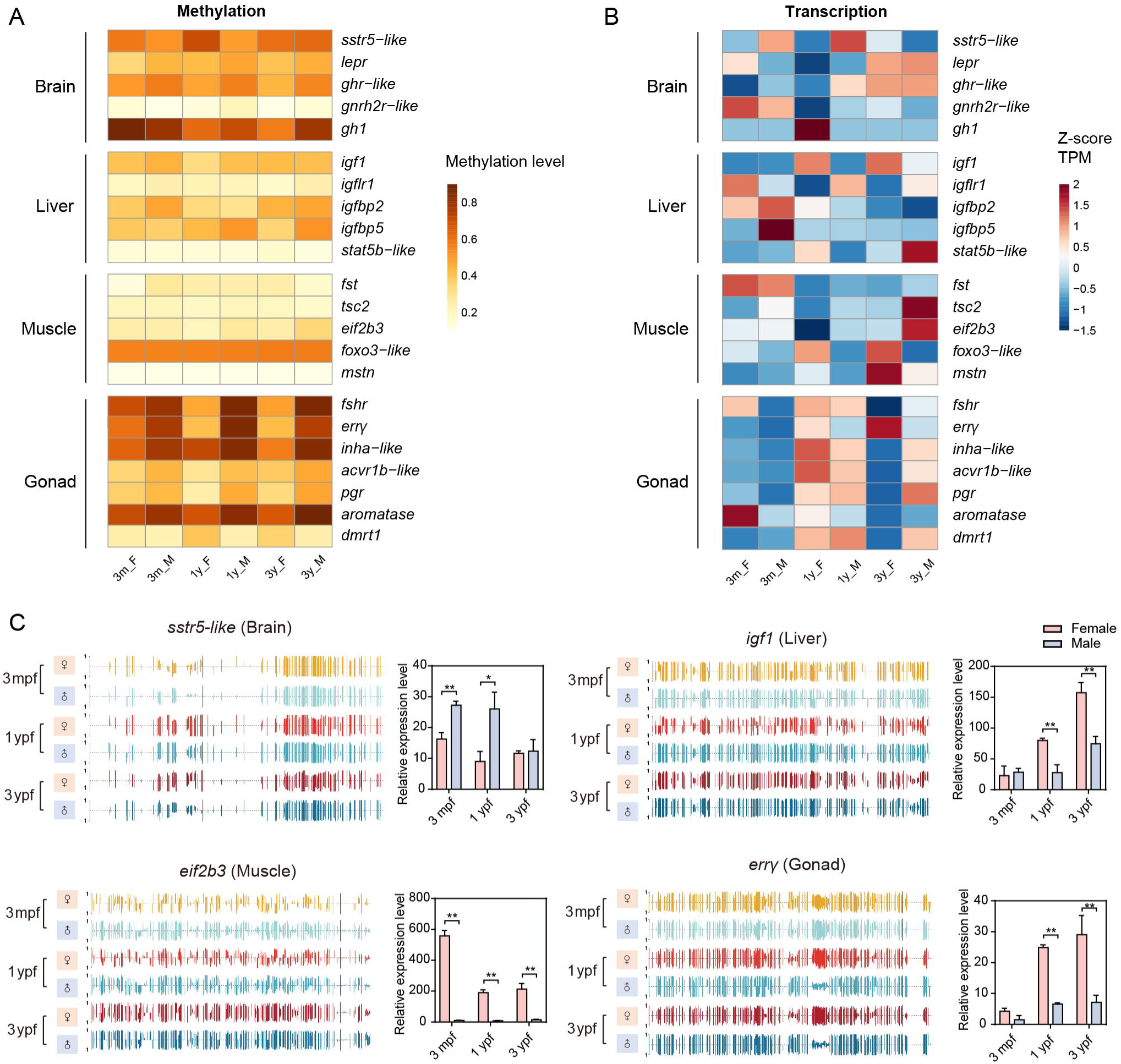
Methylation and transcription analysis of genes located on the GH/IGF network and HPG axis **A.** Methylation changes of representative genes of the GH/IGF network and HPG axis in each tissue along with development in females and males. **B**. Transcription changes of genes in the GH/IGF network and HPG axis of *C. semilaevis*. RNA-Seq TPM data was treated by Z-score normalization. **C**. Methylation and expression dynamics of *sstr5-like*, *igf1*, *eif2b3*, and *err*γ, respectively. Methylation profiles were tracked at promoter regions (transcription start site −2 kb to +500 bp). GH, growth hormone; IGF, insulin-like growth factor; HPG, hypothalamus-pituitary-gonadal.

On the HPG axis, in brain, the gonadotropin releasing hormone receptor gene *gnrhr2r-like* (XM_025061537.1) were hypo-methylated in females compared with males at 3 mpf (female = 0.14; male = 0.08). The expression levels of downstream gonadotropin genes, *cgba* (XM_025053711.1) and *cgbb* (NM_001294204.1) were significantly higher in 1 ypf and 3 ypf females than males (*P* < 0.001). In gonad, follicle stimulating hormone receptor *fshr* (NM_001294190.1) and progesterone receptor *pgr* (XM_008308970.3) showed decreased methylation level in females from 3 mpf to 1 ypf [*fshr*: female = 0.70 (3mpf), 0.47 (1ypf), 0.49 (3ypf); male = 0.80 (3mpf), 0.85 (1ypf), 0.85 (3ypf); *pgr*: female = 0.36 (3mpf), 0.25 (1ypf), 0.33 (3ypf); male = 0.42 (3mpf), 0.46 (1ypf), 0.47 (3ypf)]. Especially, the estrogen-related receptor gene *err*γ (XM_008313249.3) was hypo-methylated in females from 3 mpf to 3 ypf [female = 0.60 (3mpf), 0.41 (1ypf), 0.42 (3ypf); male = 0.78 (3mpf), 0.85 (1ypf), 0.75 (3ypf)], with the mRNA expression levels higher in females than in males from 1 ypf to 3 ypf (*P* < 0.01) (Figure 4C), indicating its important role in regulating both growth and reproduction. The methylation levels of male-determining gene *dmrt1* and female-related gene *aromatase* (*cyp19a1a*) all significantly differed between males and females from 1 ypf on. Overall, DNA methylation plays a key role in regulating the growth-related GH/IGF network and reproduction-related HPG axis.

### The W chromosome is the main target for sexual dimorphism of the methylation modifications

The W chromosome represents the primary genomic difference between female and male *C. semilaevis*. There were 306 and 921 protein-coding genes on the *C. semilaevis* W and Z chromosome, respectively [33]. To investigate whether methylation status of growth-related genes located on the W chromosome and other W-specific genes contribute to SSD, we performed comparative analyses of promoter methylation (transcription start site −2 kb to +500 bp) patterns between sex-chromosomes. Only protein-coding genes that showed high expression in at least one sample were used to exclude the noise of pseudogenes. Overall, W chromosome exhibited a higher methylation levels (0.4842) compared to both the Z chromosome (0.2657) and autosomes (0.2928) (*P* < 0.001, Wilcoxon test) (**Figure 5A**, **B**, Table S7).

**Figure 5.**
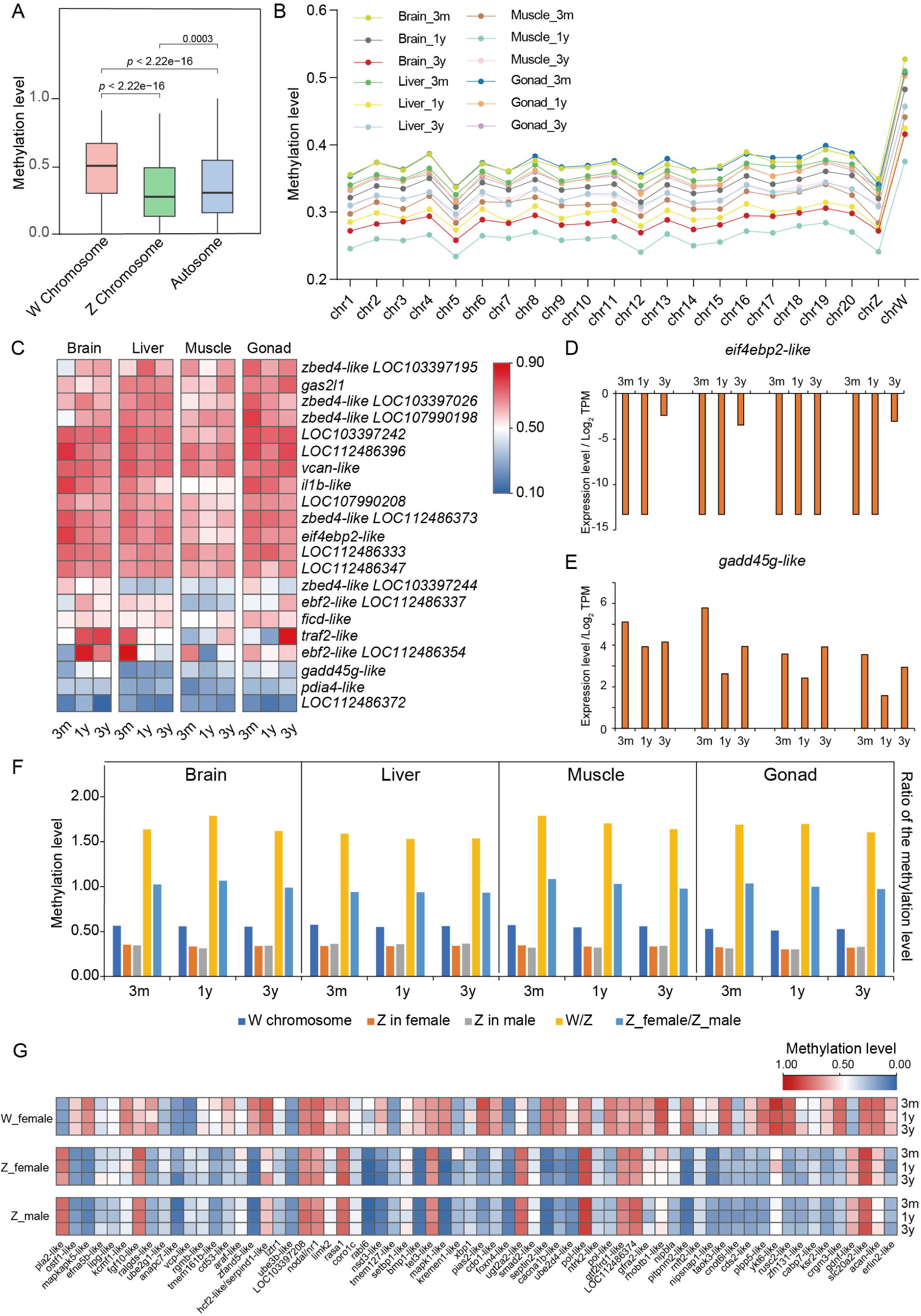
Patterns of W chromosome genes related to growth and reproduction **A.** Promoter methylation levels of different chromosomes calculating solely by protein-coding genes (data in 1 ypf female brain were used as a representative example). **B**. Chromosome methylation level of females in each stage and each tissue. **C**. Methylation profile of 21 growth- and reproduction-related genes that had no Z chromosome homolog. **D**, **E**. Relative expression levels of *eif4ebp2* and *gadd45*. Data were shown as Log_2_ TPM. **F**. Average methylation level and W/Z ratio of 66 W-genes and their Z-homologs. **G**. Methylation profile of growth- and reproduction-related genes that had homologs on the Z chromosome (data in liver were used as a representative example).

Among the 306 protein-coding genes on the W chromosome, 260 genes have homologs on the Z chromosome (Figure S9A, Table S8). On the other hand, 105 of the 306 W genes were growth- and reproduction-related, located on pathways such as “cytokines and growth factors” and “ras signaling pathway” (Figure S9B). Considering these two properties of W chromosome genes, 84 of them were shared in the two categories, and 21 growth- and reproduction-related genes were W-specific that had no homolog on the Z chromosome (Figure S9C).

Of the 21 W-specific genes related to growth and reproduction, *eif4ebp2* (eukaryotic translation initiation factor 4E-binding protein 2-like, XM_017042673.1) exhibited high methylation levels and relatively low expression levels across the life stages, especially in brain and liver (Figure 5C, D). This was consistent with numerous needs of proteins required for growth. The opposite pattern was found in the growth inhibiting gene *gadd45* (growth arrest and DNA damage-inducible protein, NM_001301180.1), which may participate in the negative feedback regulation of rapid growth in female (Figure 5C, E). Regarding the 84 shared genes, methylation level of 66 genes was detected, and their methylation state was similar between tissues (Table S9). The average methylation level on the W chromosome was higher than their Z homolog in each tissue at all life stages (average methylation ratio W_female/Z_male = 1.65), while the Z chromosome in females showed similar methylation levels to the Z chromosome in male (average methylation ratio Z_female/Z_male = 1.00) (Figure 5F). Of these genes, 43 genes on the W chromosome were more methylated than their Z homologs (W/Z > 1.2), and their expression levels were lower with a median female-to-male expression ratio of 0.25, while Z chromosome genes in females showed similar expression levels with the Z chromosome genes in male (Figure 5G, data in liver was used for example; Figure S10). Growth-inhibiting genes located on the IGF and protein degradation networks, such as *bmp1-like* (bone morphogenetic protein 1-like, XM_017042717.2), *pias2-like* (E3 SUMO-protein ligase PIAS2-like, XM_025052693.1), and *zfand5-like* (AN1-type zinc finger protein 5-like, XM_008335361.3) (Figure S9D), were correspondingly down-regulated in female than in male liver (Figure S9E). There were 7 genes less methylated than their Z homologs (W/Z < 0.8) (Figure 5G), including *pla2-like* (phospholipase A2-like, XM_008335177.3), the negative regulator of the IGF pathway (Figure S9D, E). We found that the expression level of *pla2-like* was still down-regulated on the W chromosome compared with its Z homolog (Figure S9E). The chromatin accessibility of this gene, as measured by assay for transposase-accessible chromatin-seq (preliminary data), was lower in W chromosome than its Z homolog (Figure S9F), meaning other mechanisms may also work. Overall, our results indicated that the W chromosome may participate in the regulation of SSD via its role in regulating the IGF network and protein degradation.

### Methylation is associated with rates of molecular evolution

We hypothesized that larger individuals have a faster rate of evolution of growth-related genes. Thus, we explored whether these genes evolved faster in female-biased SSD species, and how methylation functions on or along with genomic evolution. We estimated selection pressures on protein-coding genes sequence calculated as evolutionary rate (the ratio of the number of nonsynonymous substitutions per non-synonymous site relative to the number of synonymous substitutions per synonymous site; dN/dS). To do this, we used orthologous genes from fish species showing female-biased SSD (*C. semilaevis*, *S. maximus*, and *D. labrax*), male-biased SSD (*O. niloticus* and *I. punctatus*), and minor-SSD (*Oryzias latipes*, *Danio rerio* and *Oncorhynchus mykiss*) (**Figure 6A**). Minor-SSD included species where males were only slightly smaller than females.

**Figure 6.**
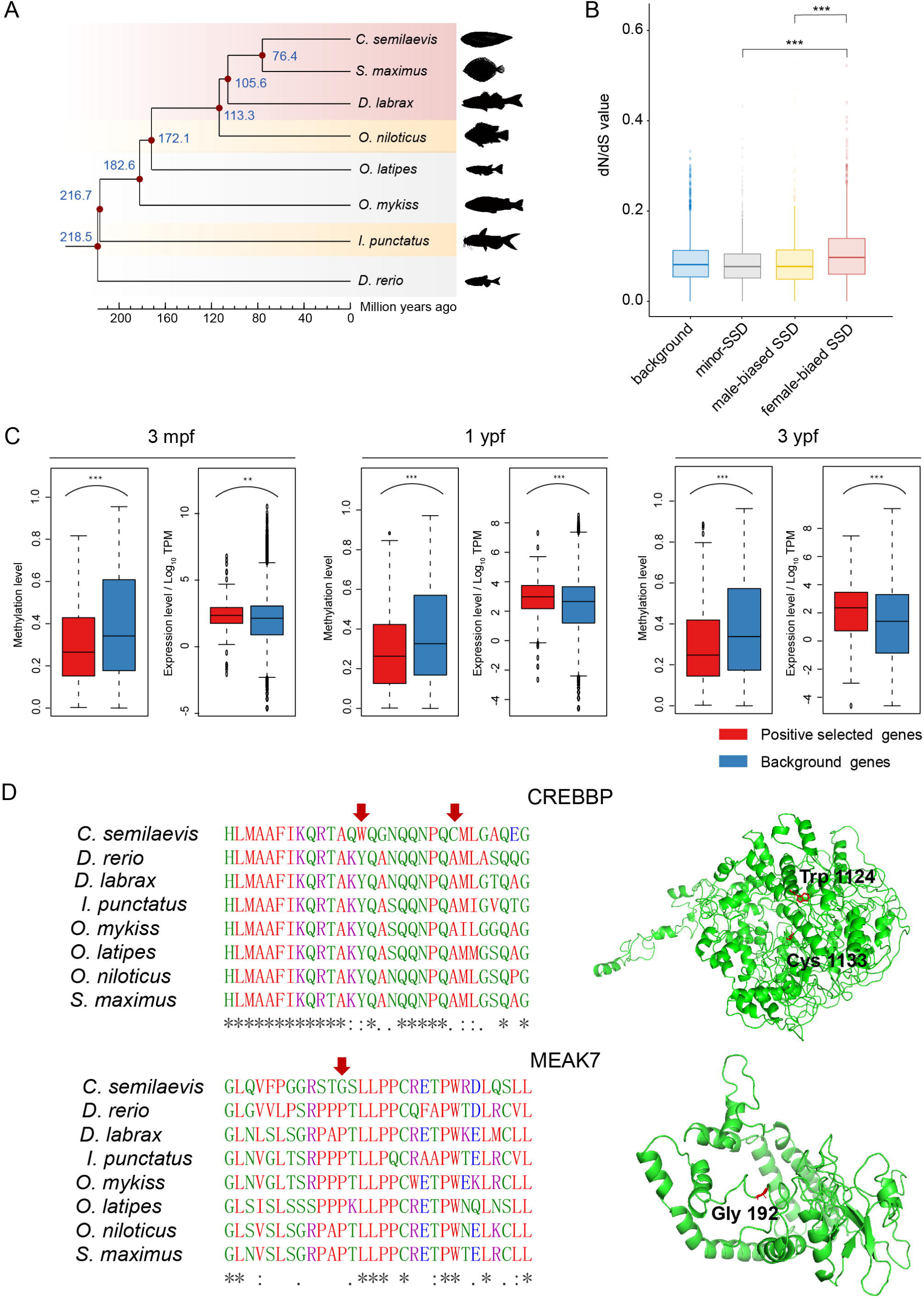
DNA methylation levels are associated with evolutionary rates **A**. Maximum-likelihood tree from teleost species used in dN/dS analysis. Branches with female-biased SSD, male-biased SSD and minor-SSD fish species are boxed in purple, yellow, and grey, respectively. Simplified species names are labeled besides the branches. **B**. Box plot of dN/dS ratios of 1066 single copy orthologous genes with higher expression in females than in males in at least one comparison from all tissues and ages. The dN/dS ratios were calculated on the branches of female-biased SSD species, male-biased SSD species, and minor-SSD species. **C**. Methylation levels and expression levels of faster evolution genes and background genes. A total of 193 genes had faster evolutionary rate specifically in *C. semilaevis* branch were selected, and compared with all other genes of *C. semilaevis*. Data in female gonad were used as a representative example. **D**. Fast-evolving sites of CREBBP and MEAK7. Left panels showed mutations in the CREBBP and MEAK7 amino acid sequences, respectively. Right panels showed the location of unique mutations in three-dimensional models of CREBBP and MEAK7, respectively. In CREBBP, the fast-evolving sites Trp 1124 and Cys 1133 were located in the CBP/p300-type histone acetyltransferase (HAT) domain. In MEAK7, the fast-evolving site (Gly 192) is on the random coil. **, *P* < 0.01, ***, *P* < 0.001, *t*-test.

A total of 2259 single-copy genes have orthologues among these eight species. Of these, 1066 genes exhibited up-regulated expression in females in at least one comparison from all tissues and ages in *C. semilaevis*. These genes showed significantly higher dN/dS ratios in branches with female-biased SSD species than in male-biased SSD and minor-SSD species (*P* < 0.001, *t*-test, Figure 6B). To investigate functional relevance, we focused on 68 growth- and translation-related genes identified through KEGG enrichment. This subset similarly displayed significantly higher dN/dS ratios in female-biased SSD species (*P* < 0.001, *t*-test, Figure S11A). Furthermore, of these 2259 genes there were 1280 female hypo-methylated DMGs across multiple tissues and developmental stages. Their evolutionary rate pattern was consistent with that of female-up-regulated DEGs, with significantly higher dN/dS ratios in female-biased SSD lineages versus male-biased/minor-SSD lineages (*P* < 0.001, *t*-test, Figure S11B).

On the other hand, of the 2259 genes there were 193 genes had faster evolutionary rate in *C. semilaevis* branch compared to other fish branches found by branch-site model (Table S10). GO enrichment analysis revealed these 193 genes were significantly overrepresented in terms including “positive regulation of cytokine production”, “nucleic acid metabolism”, and “double-strand break repair”. KEGG enrichment analysis found the “gonadotropin releasing hormone (GnRH) secretion pathway” was significantly overrepresented (Figure S12). When compared to all remaining genes, these genes were found to be less methylated and higher expressed on average than the remaining genes in all tissues (*P* < 0.05, *t*-test), except in brain where the expression level had no significant difference (**Figure 6C**, data in female gonad were used as a representative example). We also found that rapid evolution of bone development related genes CREBBP (CREB-binding protein, XP_024919985.1) and mTOR signaling activator MEAK7 (MTOR associated protein, eak-7 homolog, XP_024912177.1) were specific to the *C. semilaevis* lineage (as shown by an elevated ratio of dN/dS rate) (**Figure 6D**). These results showed that methylation levels and associated gene expression changes are more pronounced in fast-evolving genes, thereby suggesting an important interplay of DNA methylation with gene sequence involved in SSD during evolution.

## Discussion

In this study, we set out to describe the methylome and transcriptome of *C. semilaevis*, covering three life stages and four tissues with key roles in growth and development in female and male, to provide insight into the underlying methylation mechanisms responsible for the appearance of SSD. To the best of our knowledge, our study provides the first comprehensive multi-tissue and multi-stage, i.e., spatiotemporal methylation characterization to examine SSD in vertebrates.

### Role of methylation regulation in sexual size dimorphism

It has been reported that methylation may influence an organism’s phenotype according to sex and life experiences, with DNA methylation changes correlating with major life events [35−37]. Previous study in mature *C. semilaevis* (1.5 ypf) muscle and gonad also identified higher methylation levels in males and pseudomales than females, and found female hypo-methylated cell cycle-related genes as well as males and pseudomales hypo-methylated hippo signalling pathway genes associated with female-biased SSD [38]. Considering the complexity of the mechanisms involved in growth regulation and development, the phenomenon of SSD should be studied across multiple tissues and developmental stages. In the present study, we found that DNA methylation levels of males and females were similar at 3 mpf, and then became progressively different with the number of DMRs increasing across the lifetime. In general, the sex difference was not significant in all but the gonad tissues. We thought it might be because the brain, liver, and muscle tissues are responsible for a wide range of vital life processes in addition to SSD. Thus, the SSD may involve certain genes rather than a global pattern throughout the genome in these tissues. By defining the SSD specific DMGs/DEGs, the significant enrichment of KEGG pathways (*e.g.*, ribosome biogenesis, DNA replication, spliceosome, ribosome biogenesis in eukaryotes, nucleotide excision repair, and PI3K-Akt signaling) in female hypo-methylated/up-regulated genes (Figure 3C), reflected a multi-dimensional strategy to meet increasing metabolic and proliferative demands, from nucleic acid amplification to structural maintenance. One of the big questions is whether methylation changes precede or, in contrast, are the result of phenotypic changes. In our present results: (1) pathways of “development and regeneration”, “signal transduction”, “GnRH secretion”, “GnRH signaling pathway”, and “growth hormone synthesis, secretion and action” were significantly enriched from 3 mpf on in the brain and gonad (Figure 2D); (2) Many important growth-/reproduction-related genes such as IGF receptor gene *igf1r1*, IGF binding protein gene *igfbp2*, myostatin inhibiting gene *fst*, gonadotropin releasing hormone receptor gene *gnrhr2r-like*, and estrogen-related receptor gene *err*γ were differentially methylated between sexes at 3 mpf before the SSD emerge; (3) The average promoter methylation level of W chromosome genes was consistently higher than their homologs on the Z chromosome, regardless of tissue types and life stages. Taken together, these results may reflect the beginning of methyl-regulation prior to the appearance of phenotypic differences, thus suggesting that methylation changes occur first. However, whether there is a cause-consequence relationship still requires further exploration.

### Methylation regulation occurred on the genes at the top of the GH/IGF and HPG axes

GH is produced mainly by the somatotroph cells in the anterior hypophysis, induced by GH-releasing hormones and inhibited by somatotropin release inhibiting factor hormone (somatostatin) [39]. After being released and entering the blood, the GH receptor in the liver can bind GH, further activating the IGF network to regulate growth [40]. IGF network plays a major role in increasing larval body mass in *Drosophila*, and IGF-1 is critical in pre- and postnatal body growth from fish to mammals [41]. The effects of IGF-1 are mediated mainly by its binding proteins, IGFBPs, which then regulate muscle mass increases via the Akt/PKB pathway [42]. Genes associated with muscle contraction and insulin signalling were found to be under methylation regulation during skeletal muscle development in mammals [43]. In our study, genes at the top of the cascades of the GH/IGF network (from *gh* to *igfbps*) exhibited sex-related methylation differences. However, in the effector organ muscle, many selected growth-related genes did not show significant methylation differences between sexes (Figure 4B). This suggests that methylation is involved in the regulation of genes at the top of the cascades (Figure 7), which then in turn regulate their downstream effectors directly, in the case of transcription factors, or indirectly via mechanisms other than DNA methylation.

**Figure 7.**
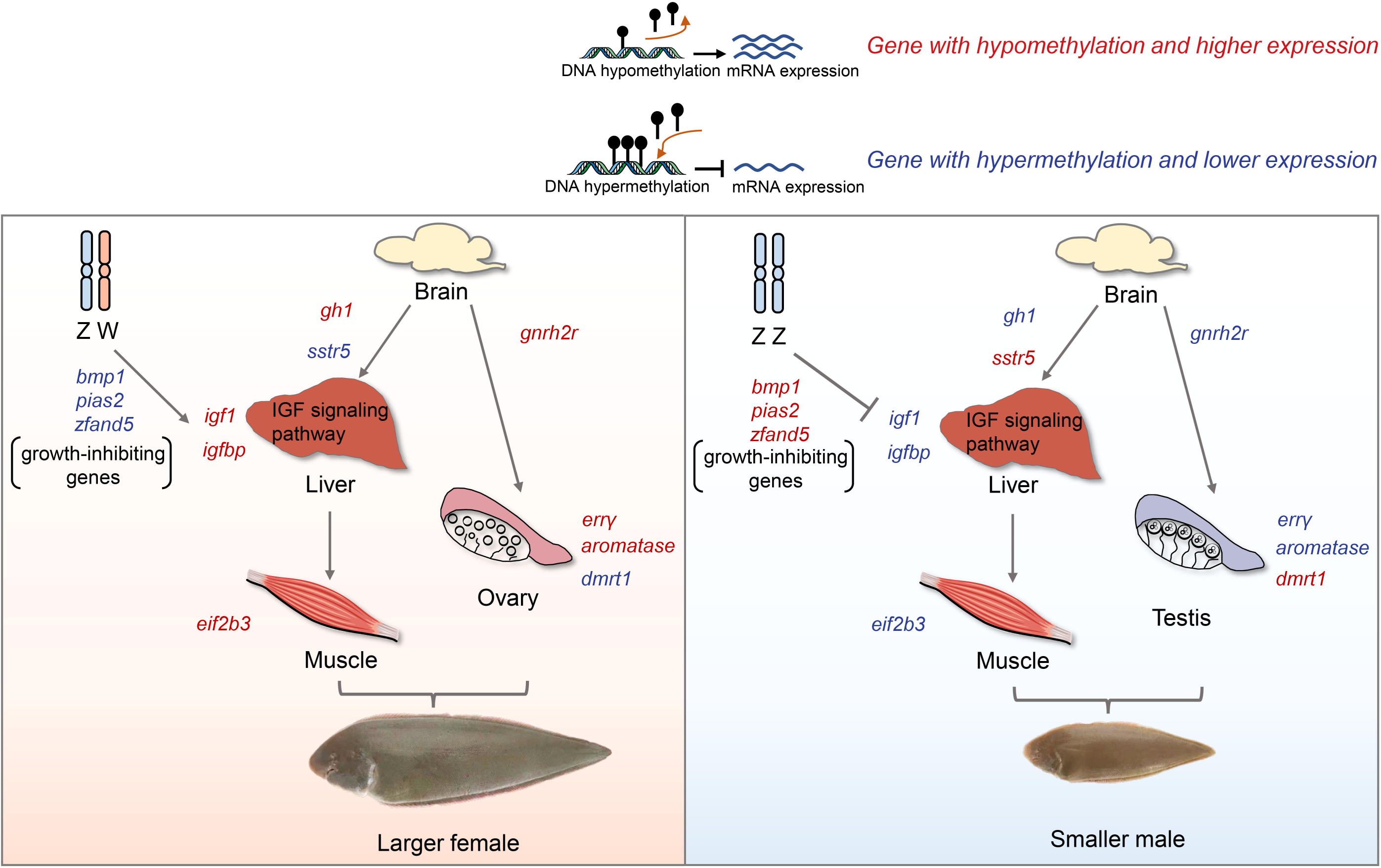
Schematic illustration proposed the role of DNA methylation on SSD regulation In female *C. semilaevis*, growth-promoting genes including the growth hormone gene *gh1* in brain, the insulin-like growth factor gene *igf1* and its binding protein gene *igfbp* in liver, and the eukaryotic translation initiation factor gene *eif2b3* in muscle, were hypo-methylated with higher expression level compared to males. While the growth-inhibiting somatostatin receptor gene *sstr5* was hyper-methylated with lower expression level in female brain. The reproductive related genes gonadotropin-releasing hormone receptor *gnrh2r*, estrogen-related receptor gene *err*γ and female-related gene *aromatase* were hypo-methylated and higher expressed in females, while the male-determining gene *dmrt1* was hyper-methylated and lower expressed in females compared with males. In addition, growth-inhibiting genes involved in the IGF and protein degradation networks, including *bmp1*, *pias2* and *zfand5*, were more methylated on the W chromosome than their Z homologues, with correspondingly lower expression levels in females than in males. Generally, key pathways and genes related to growth and reproduction can be influenced by DNA methylation, while the W chromosome may also affect the IGF network through DNA methylation. Together, they have an methylation role in regulating SSD.

On the other hand, the HPG axis plays an important role in reproduction. It begins with the release of gonadotropin-releasing hormone (GnRH), which stimulates the secretion of the gonadotropins, luteinizing hormone (LH) and follicle-stimulating hormone (FSH). These hormones act on the gonads to stimulate the production of gametes and promote gonadal release of sex steroids, thus controlling reproduction [44,45]. Methylation mechanisms were found to participate in the crosstalk between growth factors and steroid hormone [46]. In our study, the “cytokine-cytokine receptor interaction” and “environmental information processing” pathways in brain, as well as “mitochondrial biogenesis” and “ribosome biogenesis in eukaryotes” pathways in gonad were enriched from 1 ypf to 3 ypf (Figure 3D), which may be consistent with the physiological stage at which growth tends to be stable and reproduction becomes the main task. Also, we found GnRH receptor *gnrh2r-like* and estrogen-related receptor gene *err*γ were hypo-methylated in females compared with males from 3mpf to 3ypf. The methylation pattern of male-related genes *dmrt1* and female-related gene *aromatase* all showed significant differences between males and females from 1ypf on (Figure 4, Figure 7). Estrogen receptors play an important role in growth and reproduction, and are involved in the process of aromatase induced ovarian differentiation. Differences in methylation levels of sex-determining genes occur later than those of estrogen receptor genes, may indicate that sex-determining genes are associated with sexually dimorphic pattern in the presence of estrogen receptors. The situation of few genes the expression of which correlates with methylation also typically found in lots of studies nowadays [47]. Taken together, our results suggest that the upstream methylation regulation influences the transition from growth to reproduction, and may be a more economical and flexible way for animals to regulate phenotypic traits.

### The role of sex chromosomes in sexual dimorphism

The SSD phenotypes often have complex genetic architectures, although the genome of males and females is essentially the same [48]. It has been argued that sex differences in gene expression and tissue phenotype must occur downstream of the sex chromosomes that are unequally represented in males and females [49]. The sex chromosome system of *C. semilaevis* is ZZ/ZW, with recombination suppression being a prominent shared feature with the evolutionarily older avian and mammalian sex chromosomes. In *C. semilaevis*, this suppression extends across most of the chromosome. The W chromosome exhibited a higher content of pseudogenes (19.74%) compared to both the Z chromosome (3.54%) and autosomes (2.48%) [33]. Our previous study illustrated that the W chromosome has relatively higher methylation levels than autosomes in females, which probably associates with the high repeat content [34]. Considering the impact of recombination suppression and pseudogenization [50], the current study exclusively analyzed protein-coding genes, demonstrating that the proximal promoter region (PPR) of W chromosome genes also had significantly higher methylation levels than those of genes on other chromosomes, particularly their corresponding Z chromosome (Figure 5A). In humans, X-chromosome-specific hypo-methylation was found played a regulatory role in fetuses that exhibited notable sex differences in growth rate, particularly in regulating trophoblast differentiation and energy metabolism. Besides, the methylation level of the maternal X chromosome was higher than that of the paternal X chromosome [51]. Thus, the higher W-chromosome methylation level observed in this study, which correlated with SSD, may indicate that while sex chromosome systems differ, the role of methylation in regulating sex-biased gene expression could be broadly relevant.

We further focused on the growth-related genes on the W chromosome. Approximately half (43/66) of the genes were more methylated than their Z chromosome homologs. Among them, we found key growth-inhibiting genes such as *bmp1-like*, *pias2-like*, and *zfand5-like*. *bmp1* plays dual roles in growth: it can activate growth inhibiting factors such as *myostatin* by cleaving extracellular antagonists [52]; on the other hand, *bmp1* can cleave *igfbp3*, resulting in markedly reduced ability to bind IGF and thus blocked IGF-induced cell signaling in human and mouse [53]. *pias2* is an inhibitor of *stat2*, which is known to be required for myogenic differentiation [54]. *stat2* can regulate the IGF/PI3K/Akt pathway by affecting *igf2* and *hgf* expression, and knockdown of *stat2* can consistently inhibit myogenic differentiation in mammals [55]. *zfand5* participates in stimulating overall protein breakdown by the ubiquitin proteasome pathway. Lacking of *zfand5* can lead to lower rates of protein degradation and proteasomal activity, and decreased muscle atrophy in mice [56]. We also found *pla2-like*, the negative regulator of the IGF pathway [57], was significantly down-regulated on the W chromosome compared to its Z homolog. Our results indicate that the W chromosome, which is directly involved in sex determination, may also indirectly regulate sexual dimorphism phenotypes by regulating autosomal genes, mainly through IGF network regulators and protein degradation related genes (Figure 7).

### DNA methylation is associated with the process of natural selection on SSD

Previous studies in chickens and humans have shown altered CpG methylation in regions under selective pressure [58]. Thus, DNA methylation exhibited a potential adaptive and evolutionary role. To test the hypothesis that growth and reproduction have been important targets of selection, we explored evolutionary analysis of these genes in the Chinese tongue sole. Our results showed that female hypo-methylated genes, female up-regulated genes, as well as growth and translation related genes, had faster evolutionary rates in female-biased SSD species than in male-biased and minor-SSD species, which indicates that they were under higher evolutionary selective pressure. The difference between dN/dS ratios in these branches suggested that different genes may be involved in growth regulation in different species and SSD types. When focusing on the 193 faster-evolving genes in female-biased SSD *C. semilaevis* branch compared to other fish branches, the DNA methylation level of these genes was lower than that of background genes. Genes associated with size traits appear to evolve more rapidly in female-biased SSD fish, even if strong evidence of positive selection on the genes cannot be deduced since the dN/dS values are not close to 1. Thus, most of the loci in the genomes of these fish species were under purifying selection, as their dN/dS ratios were less than 1. However, female-biased SSD fish (especially their growth-related genes) may be subject to slightly stronger natural selection than other species, resulting in the retention of more nonsynonymous mutations.

We also found that CREBBP and MEAK7 were specific to the *C. semilaevis* lineage. CREBBP had two nonsynonymous mutations located on its CBP/p300-type histone acetyltransferase (HAT) domain, with the substitution at position 1124 from Tyr to Trp may affect protein function with a SIFT (sorting intolerant from tolerant) score of 0.04. CREBBP acts as transcriptional coactivator in many developmental processes, and also showed nonsynonymous mutations in giraffe (*Giraffa camelopardalis*) compared with other ruminants in an analysis of body size evolution [59]. Despite non-identical residues, recurrent HAT domain substitutions and faster evolutionary rate of this gene and across taxa may highlight its potential conserved role related to enlarged body structures. MEAK7 is known to activate alternative mTOR signaling by repressing EIF4EBP, thus regulating cell proliferation and migration [60]. *eif4ebp3,* which inhibits protein synthesis, exhibited down-regulation in females in the present study. Thus, nonsynonymous mutations of MEAK7 may be related to the larger body size in female *C. semilaevis* because of its role as an activator of the developmental pathway. Taken together, our findings provide evidence of natural selection acting on methylation variation associated with SSD traits, thus widening our understanding of the molecular basis of SSD.

## Conclusion

In conclusion, our results provide a detailed molecular picture of methyl-regulated functions in SSD. Methylation changes in the upper hierarchy of the GH/IGF and HPG axes were related to concomitant changes in gene expression. The female-specific W chromosome had higher methylation levels than autosomes and the Z chromosome, and negative regulators of the IGF pathway were more methylated than their Z chromosome homologs. Furthermore, methylation exhibits a correlation with rates of molecular evolution, thereby suggesting an important role for DNA methylation in SSD evolution. Our results contribute to understand how methylation regulation can help living organisms to allocate the resources influencing growth and reproduction between different sexes to maximize individual fitness, and to cope with the pressures of natural selection.

## Materials and methods

### Sample collection

A total of 216 healthy female and male Chinese tongue sole at the juvenile stage (3 mpf, the sex-determination program was accomplished), male maturing stage (1 ypf), and female maturing stage (3 ypf) were selected randomly from 9 tanks (three for each stage) from Laizhou Mingbo, Co (Yantai, China). Fish were fed twice a day with commercial pellets, routinely reared in filtered seawater at 22°C, and sampled at the same time to exclude differences due to seasonality. The genetic sex of each fish was identified by a transformed simple sequence repeat (SSR) primer pair as previously described [31]. The number of fish per stage, tissue and sex was nine. After body length and body weight measurement, three fish per stage, tissue and sex were used for WGBS and transcriptome sequencing. Fishes were anesthetized with 0.2% tricaine methanesulfonate (MS-222) and then sacrificed. Tissues of whole brain, liver, muscle, and gonad were sampled, divide equally and immediately stored in liquid nitrogen for simultaneous DNA and RNA extractions.

### DNA and RNA extraction

Genomic DNA of each sample was isolated with E.Z.N.A.® Tissue DNA Kit according to the manufacturer’s instructions and quantified by using a TBS-380 fluorometer (Turner BioSystems Inc., Sunnyvale, CA). High quality DNA samples (OD260/280 = ∼1.8–2.0, > 6 µg) were used to construct the WGBS libraries. Total RNA was extracted from each sample via Trizol Reagent (Invitrogen, Carlsbad, CA, USA). Then, the total RNA was quantified by Nano Drop and quality controlled by Agilent 2100 bioanalyzer (Thermo Fisher Scientific, MA, USA).

### Library preparation, sequencing for WGBS and RNA-seq

For WGBS, before bisulfite treatment, 25 ng lambda-DNA were added to the 5 μg genomic DNA for 72 samples. Then the mixed DNA was fragmented with a sonicator (Sonics &Materials) to 450 bp. After blunt ending and 3’-end addition of dA, Illumina methylated adapters were added according to the manufacturer’s instructions of the TruSeq Nano DNA LT Sample Prep Kit (Illumina, San Diego, CA, USA). Bisulfite conversion of DNA was carried out using ZYMO EZ DNA Methylation-Gold kit (ZYMO, Irvine, CA, USA) and libraries were then amplified by 12 cycles of PCR with KAPA HiFi HotStart Uracil+ReadyMix PCR Kit (KAPA, Woburn, MA, USA). Ultra-high-throughput paired-end sequencing (350 bp) was carried out using the Illumina Hiseq according to the manufacturer’s instructions. Raw Hiseq sequencing data of 72 libraries were processed by Illumina base-calling pipeline (SolexaPipeline-1.0).

For RNA-seq, the cDNA library was constructed using TruSeq RNA sample prep kit (Illumina, San Diego, CA, USA) according to the manufacturer’s instructions. After quality control, a total of 72 cDNA libraries were constructed and paired-end 100 bp reads were generated on BGI-seq 500.

### Bioinformatic analysis of WGBS and RNA-seq data

The raw paired-end reads from WGBS were trimmed and quality controlled by the Fastp software with default cutoff Q20 for Phred scores [61]. All clean WGBS reads were mapped to the reference *C. semilaevis* genome (assembly Cse_v1.0) with the BSMAP aligner (version v2.90) with default parameters [62]. Uniquely mapped reads were used to determine the cytosine methylation levels as previously stated [34]. The error rate of each library (sum of the non-conversion rate and T/C sequencing errors) was calculated as the total number of sequenced Cs divided by the total sequencing depth for sites corresponding to Cs in the Lambda genome. CpG sites with coverage ≥ 4 were selected and used to perform Principal Component Analysis (PCA) in order to detect DNA methylation patterns in different tissues across developmental stages and sexes. To distinguish true mCs from false positives, we used a model based on the binomial distribution B(n,p) following the method of Bonasio et al. [63], and only the Cs with coverage ≥ 4, mCs coverage ≥ 4, and false discovery rate (FDR) adjusted *P*-values < 0.01 were considered true positives [64].

Using the raw data from RNA-seq, adapters were removed and low-quality reads (low-quality base ratio >20%, or unknown base ratio >5%) were filtered out by SOAPnuke (version 1.4.0) [65]. Subsequently, the filtered reads of each sample were mapped to the *C*. *semilaevis* genome by HISAT2 (version 2.1.0) [66]. Gene expression levels were estimated using salmon-SEME (version 1.3.0) [67]. Then, gene expression levels were normalized by transcripts per million (TPM) to eliminate the influence of different gene lengths. After PCA analysis and pearson correlation test using Log_10_ (FPKM+1)-transformed expression data (calculated via a custom R script that employed the reshape2 package), samples with an inter-sample pearson correlation < 0.85 were eliminated, and a total of 66 cDNA libraries were retained for subsequent analysis (Table S11). Differential gene expression was identified using DESeq2 (version 1.12.3) with thresholds set as fold change of female: male > |2| and FDR-adjusted *P*-values < 0.05 [68].

### Identification of DMRs and DMGs

DMRs between females and males were detected by the DSS R package (version 2.38.0) with smoothing mode based on the CpG sites [69−71]. Differentially methylated loci (DML) were firstly identified with default parameters (*P*-value < 1e-5), then DMRs were called based on the DML requiring 50 bp minimum length and 3 minimum DML per region. Neighboring DMRs were combined if the distance was less than 100 bp. DMGs were defined as genes containing DMRs in their putative promoter regions (TSS −2 kb to +500 bp), containing at least five CpG sites that covered by at least four reads.

### Gene Ontology terms and Kyoto Encyclopedia of Genes and Genome analysis

GO (Gene Ontology) terms and KEGG (Kyoto Encyclopedia of Genes and Genome) pathways enrichment analysis was performed by using Phyper in R. Fisher’s exact test and chi-squared tests were used to estimate the enrichment and *P*-values < 0.05 was set as significance threshold. During the integrate KEGG analysis of sex-related DMGs/DEGs across different stages, the selection criteria for pathways was as follows: Step 1, KEGG enrichment analyses were conducted on DEGs between females and males from all tissues (brain, liver, muscle, gonad) and developmental stages (3 mpf, 1 ypf, 3 ypf) separately, generating 12 distinct KEGG datasets. The significance threshold was sea as *P* < 0.05. For groups yielding more than 20 enriched pathways, the top 20 most significant pathways were retained. Step 2, Within the same tissue, redundant KEGG shared across 2-3 stages were prioritized for retention, and stage-specific KEGG were sorted by *P*-values with smaller *P*-values retained preferentially. Step 3, The same analysis was performed on the duplicate and specific KEGGs across the four tissues. Finally, 30 KEGGs were retained. For each KEGG, the *P*-value of each group was listed.

### Correlation analysis between methylation and gene expression

The correlation of methylation levels of gene promoter region and gene expression was performed by using R package hexbin (version 1.28.2). The expression data were normalized as Log_10_ (TPM+0.01). Differences among means of methylation and expression data were tested by ANOVA with a Tukey HSD post-hoc test (*P*-value < 0.05).

### Comparable methylation analysis of W chromosome

The R script was used to load and merge methylation data from Z, W, and autosomes. The ggplot2 package was employed to create boxplots showing each group’s distribution of methylation levels. Wilcoxon rank-sum tests were performed to assess statistical differences among the groups, and the resulting *P*-values were displayed on the plots.

### Multiple alignment construction and evolutionary rate analyses

Eight fish species including female-biased SSD fish (*C. semilaevis*, *S. maximus*, and *D. labrax*), male-biased SSD fish (*O. niloticus* and *I. punctatus*), and minor-SSD fish (*O. latipes*, *D. rerio*, and *O. mykiss*) were used to construct phylogenetic branching and divergence time. Protein datasets were obtained from Ensembl-FTP release-103 (*S. maximus*, *D. labrax*, *O. niloticus*, *I. punctatus*, *O. latipes*, *D. rerio*, and *O. mykiss*) and GCF_000523025.1 (*C. semilaevis*). Evolutionary rate (the ratio of the number of nonsynonymous substitutions per non-synonymous site relative to the number of synonymous substitutions per synonymous site, dN/dS) was analyzed by using orthologous genes and calculated using the PAML [72]. The orthologous genes with higher expression in female *C. semilaevis* in at least one comparison from all tissues and ages were retrieved. There are three models of PAML. The branch models allow the dN/dS ratios to vary among branches in the phylogeny and are useful for detecting evolutionary rate on particular lineages. The site models allow the dN/dS ratio to vary among codons or amino acids in the protein. The branch-site models aim to detect positive selection that affects only a few sites on prespecified lineages. To detect if lineage underwent fast evolution, we chose the branch model to detect the evolution rate and dN/dS ratios on the branches were calculated using the two-ratio branch model. The female-biased SSD branch was set as foreground branch (model 2), and background branch was set as model 1. We also chose this model according to the evolutionary analyses of body size in ruminants [59]. The 193 genes with faster evolutionary rate in *C. semilaevis* branch were analyzed by branch-site model. Student’s *t*-test was used to test for pairwise differences in dN/dS between the 4 groups.

### qRT-PCR validation

Candidate DEGs from each tissue participating in growth- and reproduction-associated pathways (*sstr5-like* in the brain, *igf1* in the liver, *eif2b3* in the muscle, and *err*γ in the gonad) were selected to validate the RNA-seq data by qRT-PCR (Table S12). β*-actin* gene was used as the internal control. One microgram total RNAs for high-throughput transcriptome sequencing was reverse transcribed into cDNA with the PrimeScript™ RT reagent Kit with gDNA Eraser (Takara, Japan). Then, qRT-PCR was performed using QuantiNova™ SYBR Green PCR Kit (Qiagen, Germany) in 20 μL reactions, containing 10 μL SYBR Green PCR Master Mix (2×), 2 μL QN ROX Reference Dye, 0.7 μM forward primer, 0.7 μM reverse primer, and 1 μL cDNA. The cycling program was carried out at 95°C for 2 min, followed by 40 cycles of 95°C for 5 s and 60°C for 10 s; this was followed by a melting curve analysis in an ABI StepOnePlus Real-Time PCR system (Applied Biossystems, USA). Reactions were performed in triplicate. The relative expression fold changes of these genes were analyzed using the 2^−ΔΔCt^ method.

## Ethical statement

All work involving Chinese tongue sole was carried out according to the ethical principles of animal welfare of the Yellow Sea Fisheries Research Institute, Chinese Academy of Fishery Sciences (Qingdao, China). All animal experiments were approved by the Institutional Animal Care and Use Committee (IACUC) of YSFRI, CAFS.

## Authors’ contributions

**QW**: Conceptualization, Investigation, Writing - Original Draft. **XH**: Sofware, Data curation, Writing– original draft. **DA**: Validation. **KL**: Methodology, Data curation. S**L**: Data curation. **BF**: Resources, Investigation. **RW**: Resources, Validation. **MR:** Writing– original draft, Writing– review & editing. **YL**: Resources, Validation. **HW**: Validation. **LT**: Validation. **FP**: Writing– original draft, Writing– review & editing. **CS**: Supervision, Project administration, Funding acquisition. All authors have read and approved the final manuscript.

## Competing interests

The authors have declared no competing interests.

## Data Access

All raw data of the methylome and transcriptome are deposited in the National Center for Biotechnology Information (NCBI) database (BioProject no. PRJNA724919).

## Supplementary material

**Figure S1** DNA methylation of *C. semilaevis* **A**. Average body length and body weight of *C. semilaevis*, according to sex and age (n = 9). **B**. Percentage of mCs in the mCG, mCHG, and mCHH contexts in *C. semilaevis*. **C**. DNA methylation levels in whole genome and different genomic regions including exon, gene, intergenic, intron and promoter. The color gradient from red to green showed hyper-methylation and hypo-methylation, respectively. **D**. DNA methylation levels in functional regions of gene. The gene features included the upstream, first exon, first intron, internal exon, internal intron, last exon and downstream. Data from the brain as a representative example. ***, *P* < 0.001, *t*-test.

**Figure S2** Integrated analysis of the genome-wide DNA methylation profiles **A**. PCA based on all methylation pattern of all genes in 72 samples. Samples were clustered by different tissues. **B**. Methylation levels of special genomic element including LINEs, SINEs, CpG islands, and promoters in females (red) and males (green) from 3 mpf to 3 ypf. **C**. Number of DMRs between female and male in different developmental stages of four tissues. DMRs are defined as hypo-methylated in females (blue) or hyper-methylated in females (red). PCA, principal component analysis; LINE, long interspersed nuclear element; SINE, short interspersed nuclear element.

**Figure S3** Heatmap of SSD-specific female hypo-/hyper-methylated DMGs in each tissue The methylation level of each SSD-specific female hypo-methylated DMGs in brain (**A**), liver (**C**), muscle (**E**), and gonad (**G**), as well as SSD-specific female hyper-methylated DMGs in brain (**B**), liver (**D**), muscle (**F**), and gonad (**H**) were shown at each stage.

**Figure S4** KEGG enrichment analysis of SSD-specific DMGs. **A**. Venn diagram of KEGG pathways enriched by SSD-specific female hypo-methylated DMGs in four tissues. **B**. Venn diagram of KEGG pathways enriched by SSD-specific female hyper-methylated DMGs in four tissues. **C**, **E**, **G**. KEGG enrichment analysis of SSD-specific female hypo-methylated DMGs in brain (**C**), liver (**E**), and muscle (**G**). **D**, **F**, **H**. KEGG enrichment analysis of SSD-specific female hyper-methylated DMGs in brain (**D**), liver (**F**), and muscle (**H**). Pathways were selected using a significance threshold of *P* < 0.05.

**Figure S5** Relationship of DNA methylation levels of promoter regions and the relative expression levels of corresponding genes Scatterplots were used to represent the relationship. The Pearson correlation coefficients (*R*) between the promoter methylation levels and expression levels were shown at the top of each panel. The horizontal axis represented the methylation levels and the vertical axis represented the expression levels.

**Figure S6** Heatmap of SSD-specific female up-/down-regulated DEGs in each tissue The expression level of each SSD-specific female up-regulated DEGs in brain (**A**), liver (**C**), muscle (**E**), and gonad (**G**), as well as SSD-specific female down-regulated DEGs in brain (**B**), liver (**D**), muscle (**F**), and gonad (**H**) were shown at each stage.

**Figure S7** Selected genes in the GH/IGF network and HPG axis of *C. semilaevis* The assumed regulatory pathway was drawn based on published literatures.

**Figure S8** Relationship of DNA methylation levels of promoter regions and the relative expression levels of GH/IGF network and HPG axis related genes Scatterplots was used to represent the relationship. The Spearman correlation coefficients (*R*) between the promoter methylation levels and expression levels were shown at the top of the panel. The horizontal axis represented the methylation levels and the vertical axis represented the expression levels as Log_2_ (TPM + 0.01).

**Figure S9** Comprehensive analysis of the W chromosome genes **A**. Frequency distribution of protein identities for the W-Z homologs. **B**. Functional annotation of the 306 W chromosome genes as shown by the significantly enriched KEGG pathways (*P* < 0.05). **C**. Venn diagram of growth- and reproduction-related genes and genes that have homologs on the Z chromosome. **D**. Methylation level of the genes *bmp1-like*, *pias2-like*, *zfand5-like*, and *pla2-like* on the W chromosome, and their corresponding Z homologs. **E**. Expression level of W chromosome genes *bmp1-like*, *pias2-like*, *zfand5-like*, and *pla2-like*, and their corresponding Z homologs. Data were shown as Log_2_ TPM. F. Chromatin accessibility profiles of *pla2*-like on W chromosome and Z chromosome, respectively (unpublished data).

**Figure S10** Comparative expression analysis of the 43 hyper-methylated genes on the W chromosome A. Scatter plots represented the correlation between Log_2_ (TPM + 0.01) of W chromosome genes in females (W_female) and their Z chromosome homologs in males (Z_male). **B**. Scatter plots represented the correlation between Log_2_ (TPM + 0.01) of Z chromosome genes in females (Z_female) and their Z chromosome homologs in males (Z_male). **C**. Boxplot of Log_2_ (W_female TPM / Z_male TPM). Black line indicated the median, lower and upper edges standed for the 25th and 75th percentiles.

**Figure S11** The dN/dS ratios of growth- / translation-related genes and female hypo-methylated genes in female-biased SSD, male-biased SSD, and minor-SSD species Box plot of dN/dS ratios of 68 growth- / translation-related genes (**A**), and female hypo-methylated genes in at least one comparison from all tissues and ages (**B**). The dN/dS ratios were calculated on the branches of female-biased SSD species, male-biased SSD species, and minor-SSD species.

**Figure S12** GO terms classification and KEGG pathway analysis of 193 genes faster evolved in *C. semilaevis*

**Table S1** Statistic of WGBS-seq for each sample

**Table S2** Clean reads mapping information of each library

**Table S3** Number of cytosines covered in each context

**Table S4** Methylated CpGs detected in each library

**Table S5** The methylation level of each library

**Table S6** The list of 205 genes mainly located on the GH/IGF network and HPG axis based on the literature

**Table S7** Chromosome methylation level of each library

**Table S8** List of W chromosome genes possessing Z chromosome paralogs

**Table S9** Methylation level of 84 W chromosome genes possessing Z chromosome paralogs and related to growth and reproduction

**Table S10** List of genes with faster evolutionary rate specifically in *C. semilaevis* branch

**Table S11** The number of *C. semilaevis* samples used for RNA-seq in each group

**Table S12** Primers used in this study

